# Salt-enhanced surface display in the halotolerant yeast *Debaryomyces hansenii*

**DOI:** 10.64898/2026.10.07.757289

**Authors:** Antonia Paić, Tea Martinić Cezar, Béatrice Vallée, Renata Teparić, Bojan Žunar

## Abstract

The halo-, xero-, and osmotolerant oleaginous yeast *Debaryomyces hansenii* can grow in extreme environments, with some isolates tolerating up to 4 M NaCl. As such, it could serve as a robust microbial chassis for non-sterile biotechnological processes based on seawater or saline side streams. However, its biotechnological toolbox lacks key genetic tools, particularly for inducible expression, secretion, and covalent anchoring of proteins to the outer cell surface. To address this gap, we characterised how growth in a medium supplemented with 1 M NaCl remodels the *D. hansenii* cell wall. We also developed strong constitutive, stationary-phase, and NaCl-inducible expression vectors, and designed protein tags for C- and N-terminal cell wall anchoring. These tools enabled functional surface display of β-lactamase, acetylxylan esterase, and lipase A via native GPI-dependent and heterologous Pir-based anchors, while immunoblotting detected the reporters in both covalently and non-covalently bound cell wall fractions. Activities of surface-displayed enzymes increased by more than 80 % under hypersaline conditions, while DhCcw12-based C-terminal anchoring outperformed Pir-based N-terminal anchoring. Thus, we retooled the *D. hansenii* cell wall into a catalytically active biointerface and established this yeast as an osmotolerant chassis for hypersaline surface display.

**Graphical abstract:** 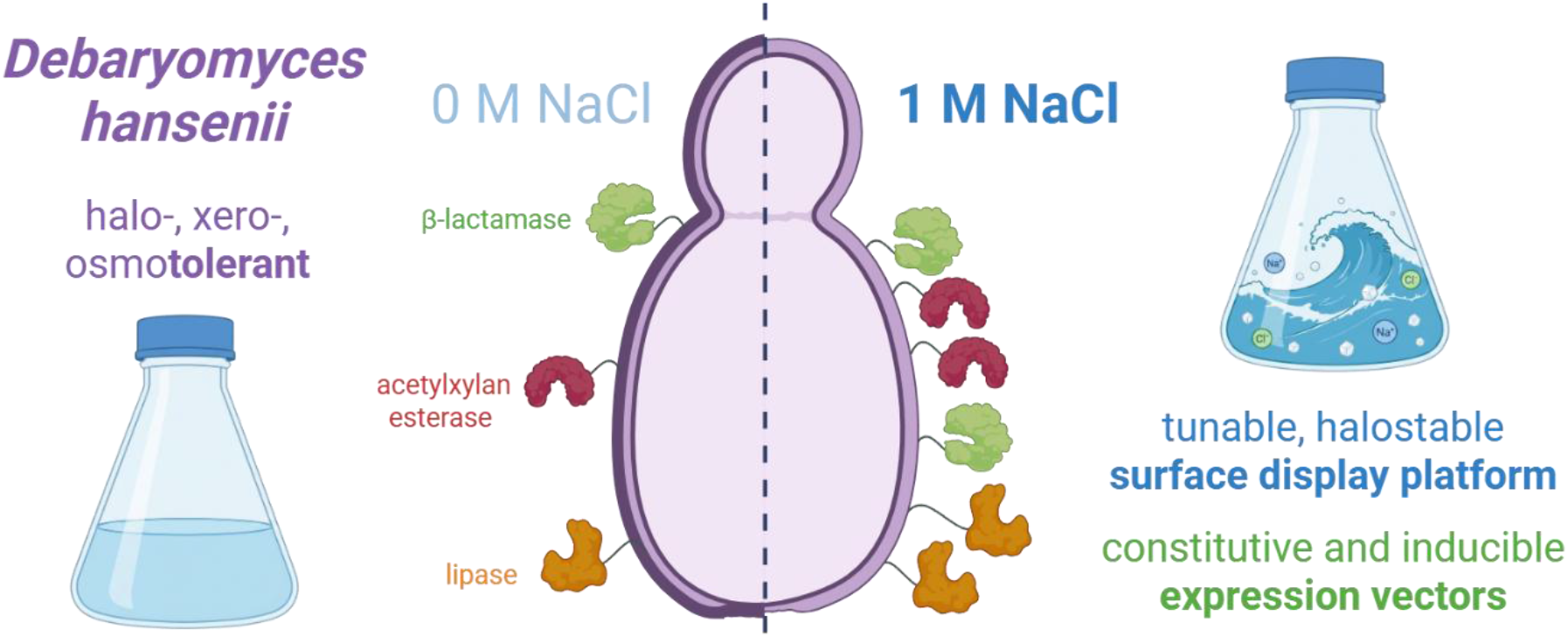

## 1. Introduction

Most microbial cell factories thrive in conditions that also favour the rapid growth of environmental contaminants, necessitating costly aseptic techniques and continually risking bioprocess failure. An alternative strategy would be to develop microbial cell factories that prefer conditions inhibitory to most microbes, thus embedding contamination control into the cultivation environment (Estrada et al., 2023a; Margesin and Schinner, 2001). Halophiles, such as the yeast *D. hansenii*, are one such group of microbes, thriving in high-salt environments (Estrada et al., 2023b; Prista et al., 2016).

*D. hansenii* is a widely distributed oleaginous ascomycetous yeast adapted to osmotic, saline, and low-water-activity environments, including seawater, saline soils, and fermented foods (Navarrete et al., 2022a). In cheese brines, it is among the most frequently detected yeasts, with its brine-derived isolates showing substantial lipolytic or proteolytic activity and producing diverse volatile profiles (Innocente et al., 2023). Selected *D. hansenii* strains also improve the aroma and sensory properties of fermented meats (Yang et al., 2024). Beyond food fermentation, this species can be used to produce riboflavin (Villarreal-Huerta et al., 2026) and xylitol (Morais et al., 2023).

*D. hansenii*’s ecological breadth relies on its unusually flexible physiology. This yeast maintains osmotic balance by controlling sodium and potassium homeostasis, accumulating glycerol, and adjusting membrane transport and metabolism (Breuer and Harms, 2006), with some isolates even growing in media supplemented with up to 4 M NaCl (Norkrans, 1966). In *D. hansenii*, NaCl and KCl elicit distinct, pH-dependent effects, with elevated sodium improving cellular performance under some conditions (Navarrete et al., 2021). Transcriptomic and phosphoproteomic analyses likewise showed that in this species, sodium and potassium trigger distinct regulatory programmes rather than a generic osmotic-stress response (Navarrete et al., 2022b), with sodium accumulation promoting growth and reshaping central carbon metabolism through the glyoxylate shunt (Ruiz-Pérez et al., 2023). Thus, *D. hansenii* physiology is adapted to a saline environment.

These properties make *D. hansenii* attractive for processes in which salinity is unavoidable or deliberately imposed. A marine strain grew efficiently in seawater-based media containing mixed sugars and inexpensive nitrogen sources, producing protein- and lipid-rich biomass with cell-associated phytase activity (Donzella et al., 2021). Moreover, *D. hansenii* producing recombinant proteins could grow in salt-rich dairy side streams (Estrada et al., 2023a), while combining dairy and biopharmaceutical side streams enabled open, non-sterile cultivation of *D. hansenii* without fresh water or additional nutrients (Estrada et al., 2023b). These studies position *D. hansenii* as a host for whole-cell biocatalysis and the production of recombinant proteins in saline environments. However, fully exploiting this potential requires *D. hansenii* expression and protein localisation systems adapted to hypersaline conditions.

Surface display is a technique that converts the cell wall’s outer surface into a catalytically active biomaterial. For this purpose, recombinant proteins are routed through the secretory pathway and tethered to the cell wall through GPI-linked proteins, agglutinins, or Pir proteins, thus combining protein production and immobilisation within a single cell (Li et al., 2024; Martinić Cezar et al., 2026). For example, *Saccharomyces cerevisiae* could surface display a multi-enzyme complex that completely depolymerised extracellular polyethylene terephthalate (Gulati and Sun, 2025), as well as a urea amidolyase that remained active through repeated treatment cycles (Wang et al., 2024). However, display performance depends strongly on the passenger protein and the selected secretion signal, fusion partner, and anchoring domain (O’Riordan et al., 2024). Thus, whether the salt-adapted cell wall of *D. hansenii* can support functional surface display, until now, remained unclear.

Here, we developed a genetic toolkit for functional surface display in *D. hansenii* under standard and hypersaline conditions. We characterised how growth in medium supplemented with 1 M NaCl remodels the cell wall, identified promoters for constitutive, stationary-phase, and NaCl-inducible expression, and adapted native DhCcw12 and heterologous ScPir2 as secretion and anchoring modules. The resulting system successfully displayed β-lactamase, acetylxylan esterase, and lipase A, whose activities increased by more than 80 % in NaCl-supplemented medium. Thus, the adaptations that allow *D. hansenii* to thrive at high salinity also support cell-surface biocatalysis under conditions poorly tolerated by conventional production hosts.

## 2. Materials and methods

### 2.1. Media and growth conditions

*Escherichia coli* was cultivated at 37 °C and 180 rpm in 2xYT liquid medium (16.0 g/l tryptone, 10.0 g/l yeast extract, 5.0 g/l NaCl) or on LB plates (10.0 g/l tryptone, 5.0 g/l yeast extract, 5.0 g/l NaCl, 15.0 g/l agar), both supplemented with 100 µg/ml ampicillin. *D. hansenii* was grown at 30 °C and 180 rpm in standard YPD medium (20.0 g/l peptone, 10.0 g/l yeast extract, 20.0 g/l glucose, and 20.0 g/l agar for solid media). Transformants were selected on media supplemented with 100 µg/ml of clonNAT (Jena Bioscience, Jena, Germany). For enzyme activity assays, flow cytometry, and confocal microscopy, cells were grown in a complete chemically defined medium (6.70 g/l Difco Yeast nitrogen base with ammonium sulphate and without amino acids, 0.77 g/l -his drop-out (cat. no. DCS 0079, Formedium, Swaffham, United Kingdom), 0.08 g/l histidine (cat. no. H8000, Sigma-Aldrich, St. Louis, United States), supplemented with 1.0 M NaCl as required). The medium was adjusted to pH 6.0 and filter-sterilised (0.22 µm), after which 20.0 g/l glucose was added separately.

To control the *DhPHO5* promoter, cells were grown overnight in chemically defined medium supplemented with 1.0 g/l KH_2_PO_4_ until reaching the stationary phase. They were then reinoculated into the same medium to 0.5 OD_600_/ml and grown for approximately 10 h until reaching the exponential phase of 2 OD_600_/ml. The cells were harvested, washed, and transferred to a phosphate-free induction medium for 16 h. This induction medium was adjusted to pH 6.0 before filter-sterilisation (0.22 µm), after which 0.4 % v/v riboflavin, 0.1 % v/v biotin, and 20.0 g/l glucose were added, as in Martinić Cezar et al. (2025).

### 2.2. Plasmid and strain construction

Plasmids were constructed in NEB Stable *E. coli* using HiFi Assembly (New England Biolabs, Frankfurt am Main, Germany), according to the manufacturer’s instructions. The plasmid sequences are provided in Supplementary Table 1. Plasmid DNA was isolated with the NucleoSpin Plasmid Mini Kit, while the PCR fragments used for transformation were extracted from an agarose gel with the NucleoSpin Gel and PCR Clean-up Kit (Macherey-Nagel, Duren, Germany). All constructs were verified by restriction digest and Sanger sequencing, with results matching the computed restriction maps. Primer synthesis was outsourced to Macrogen Europe (Amsterdam, Netherlands), gene synthesis to Integrated DNA Technologies (Leuven, Belgium), and Sanger sequencing was provided by Microsynth (Balgach, Switzerland).

*D. hansenii* JCM 1990ᵀ (Japan Collection of Microorganisms, RIKEN, Japan) was transformed by electroporation, using a MicroPulser electroporator (Bio-Rad, Hercules, United States). A fresh stationary-phase culture was inoculated into 100 ml of YPD medium to a starting 0.5 OD_600_/ml and grown at 30 °C and 180 rpm until reaching 3.0 OD_600_/ml. The cells were washed twice with 25 ml of sterile deionised water (sdH_2_O), resuspended in 20 ml of 100 mM lithium acetate containing 35 mM DTT, and incubated at 30 °C and 180 rpm for 45 min. The cells were washed in ice-cold sdH_2_O, resuspended in 250 µl of cold 1.0 M sorbitol, and split into 50 µl aliquots, each of which was mixed with 200 ng of the transforming DNA. Electroporation was performed at 1.8 kV in a 0.2 cm cuvette. Immediately after the pulse, the suspension was resuspended in 1 ml of 0.5x YPD supplemented with 1.0 M sorbitol, transferred to a 15 ml tube, and shaken at 30 °C and 180 rpm for 1.5 h before plating on selective plates.

### 2.3. Yeast staining, flow cytometry, and confocal microscopy

To examine the cell wall properties of *D. hansenii*, the yeast culture was grown in chemically defined medium with or without 1.0 M NaCl. Cells were harvested for staining in either the exponential (1.0-2.0 OD_600_/ml) or the stationary phase (after two days of growth), then washed with PBS (pH 7.0). The washed cells were resuspended in 100 µl of PBS and mixed with either 10 µl of 100 µg/ml PBS-dissolved concanavalin A-FITC (to visualise mannoproteins; C7642, Sigma-Aldrich, St. Louis, United States), 100 µl of 25 % Calcofluor White (to visualise chitin; 910090, Sigma-Aldrich, St. Louis, United States), 10 µl of 5 mg/ml PBS-dissolved Aniline Blue (to visualise β-glucans; 415049, Sigma-Aldrich, St. Louis, United States), or 10 µl of 100 µg/ml PBS-dissolved poly-L-lysine-FITC (to measure electronegativity; P3543, Sigma-Aldrich, St. Louis, United States), and staining proceeded for 30 min in the dark. After staining, 300 µl of PBS was added to the mixture, and cells were analysed by flow cytometry and confocal microscopy, as in Šupljika et al. (2025).

### 2.4. β-lactamase assay

β-Lactamase activity was quantified with a nitrocefin-based spectrophotometric assay. Nitrocefin, a cell-impermeable chromogenic cephalosporin, is hydrolysed by β-lactamase at its β-lactam ring to a red product, whose concentration was determined by measuring the absorbance at 482 nm.

The activity of the surface-displayed β-lactamase was measured as described previously (Martinić Cezar et al., 2025), using 0.25 OD_600_ of induced cells per reaction.

To assess intracellular β-lactamase activity, cells were first permeabilised with digitonin (cat. no. 407565000, Thermo Fisher Scientific, Italy) to facilitate substrate uptake. Induced cells were harvested and washed with sdH_2_O, then with MOPS buffer (100 mM, pH 7.0), by centrifuging at 3,000 rpm for 5 min. The pellet was resuspended in 10 mg/ml digitonin to 10 OD_600_/ml and incubated at 30 °C for 30 min. The cells were then washed with sdH_2_O and potassium phosphate buffer (KP buffer, 50 mM, pH 7.0) by centrifuging at 8,000 rpm for 3 min. The permeabilised cells were subjected to the nitrocefin assay as described in Martinić Cezar et al. (2025), using 0.05-0.15 OD_600_ of cells per reaction. β-Lactamase activity was calculated as A_482_ normalised to OD_600_. All measurements were performed in biological and technical triplicates.

### 2.5. Acetylxylan esterase and lipase A assay

As acetylxylan esterase and lipase A were expressed from the *DhPHO5* promoter, cells were cultivated and induced as described above, in medium with or without 1.0 M NaCl. The cells were washed in sdH_2_O and resuspended in KP buffer to 100 OD_600_/ml. For the activity assay, 20 µl of this cell suspension was mixed with 470 µl of KP buffer and 10 µl of 4-nitrophenyl acetate (900 ng/ml, dissolved first in DMSO at 9.0 µg/ml and then diluted 1:10 in water; N8130, Sigma-Aldrich, St. Louis, United States), and incubated at room temperature for 5 min. The reaction was stopped by centrifuging at 8,000 rpm for 2 min, and the absorbance of the released 4-nitrophenol was measured at 405 nm. A strain expressing neither acetylxylan esterase nor lipase A served as the blank. The assay was performed in biological and technical triplicates.

### 2.6. Isolation of proteins and immunoblotting

Yeast cells were cultivated and induced as described above. Non-covalently and covalently bound cell wall proteins were isolated following the methodology described in Lozančić et al. (2021). Briefly, cells were washed and resuspended in potassium phosphate buffer (KP8 buffer, 50 mM, pH 8), mechanically disrupted with glass beads in a BeadBug homogeniser (Benchmark Scientific, Edison, United States), transferred to a new tube, and washed four times with KP8 buffer. Non-covalently bound proteins were extracted by adding 1 ml of Laemmli buffer (6.0 mg/ml Tris, 0.76 mg/ml EDTA, 20 mg/ml SDS, 5 % v/v β-mercaptoethanol, 0.001 % v/v bromophenol blue) and boiling for 10 min, after which the supernatant was collected.

To extract covalently bound proteins, the pellet remaining after non-covalent protein extraction was again treated with 1 ml of Laemmli buffer and boiled for 10 min, after which the supernatant was discarded. Such stripped cell walls were washed in potassium phosphate buffer (50 mM, pH 6) and treated with β-glucanase (NZYtech, Lisbon, Portugal) in the same buffer at 55 °C for 3 h to release all covalently bound proteins. Alternatively, they were washed with sdH_2_O and treated with 30 mM NaOH overnight at 4 °C to release Pir-bound proteins. The extracted protein samples were standardised for loading by normalising the volume of extract used for electrophoresis to the mass of the wet cell walls measured before extraction of the non-covalently bound proteins.

To isolate secreted proteins, the cells were harvested by centrifugation, and the supernatant collected. To 22.5 ml of the supernatant, 15 ml of cold (−20 °C) acetone was added, and proteins were precipitated overnight at 4 °C. The suspension was centrifuged at 10,000 rpm and 4 °C for 60 min, and the precipitate was dissolved in 50 µl of Laemmli buffer.

Proteins were resolved on 10 % SDS-PAGE gels and transferred to a nitrocellulose membrane with the semidry Trans-Blot Turbo Transfer System (Bio-Rad, Hercules, United States), as in Novačić et al. (2021). Membranes were blocked overnight at 4 °C in blocking buffer (6.0 g/l TRIS, 8.8 g/l NaCl, 0.1 % Triton X-100, pH 7.5) with 1 % milk and probed with a high-affinity anti-HA peroxidase-conjugated antibody (1:2,500 dilution; 11667475001, Roche, Basel, Switzerland). Blots were developed with Clarity Western ECL substrate (1705061, Bio-Rad, Hercules, United States) and visualised with a C-DiGit Blot scanner (LI-COR Biosciences, Lincoln, United States).

### 2.7. Bioinformatic analyses

Published RNA-seq datasets for *D. hansenii* grown in standard medium vs. medium supplemented with 1 M NaCl (Navarrete et al., 2022b; Sánchez, 2021) and for *S. cerevisiae* BY4743 before and after 3 h of exposure to 0.7 M NaCl (GEO accession GSE226246) (Robinson et al., 2023) were reanalysed. Each dataset comprised three biological replicates per condition. Differential expression was analysed in R 4.4.2 using DESeq2 (Love et al., 2014), with NaCl treatment as the only factor. Genes with one or fewer reads across all six samples were removed prior to analysis. DESeq2 was run with default normalisation, dispersion estimation, and negative binomial modelling. Wald-test *p*-values were adjusted for multiple testing using the Benjamini-Hochberg method, and genes with an adjusted *p*-value below 0.05 were considered differentially expressed. Log_2_(fold-change) estimates for NaCl-treated relative to untreated cells were shrunk using apeglm (Zhu et al., 2019). Sample relationships were evaluated using Poisson distances calculated from the raw count matrices and by principal component analysis after variance-stabilising transformation.

Genes associated with the cell wall integrity and high-osmolarity glycerol pathways, and their downstream processes, were selected based on the KEGG pathway maps sce04011 and dha04011. *S. cerevisiae* common gene names were mapped to systematic ORF and SGD identifiers using the AnnotationDbi and org.Sc.sgd.db Bioconductor packages, whereas the corresponding *D. hansenii* genes were selected by their DEHA2 locus identifiers. Shrunken log_2_(fold-changes) were mapped to a diverging colour scale, with values outside the range −4 to +4 truncated at the respective limit.

Statistical analyses for reporter activities were performed in R 4.4.2, using a two-sided Welch’s *t*-test, which does not assume equal variances.

## 3. Results

### 3.1. Growth in NaCl-supplemented medium restructures the *D. hansenii* cell wall

To determine how hypersalinity affects the cell surface available for protein display, we first characterised the *D. hansenii* cell wall in standard medium and medium supplemented with 1 M NaCl. Supplementation with NaCl altered the cell wall composition in both the exponential and stationary phases (Figure 1A). Compared to cells grown in standard medium, those grown in NaCl-supplemented medium showed markedly lower chitin and β-glucan content and marginally lower mannoprotein content of the cell wall. The cells also differed in cell wall electronegativity and were smaller. The same trends were evident in the fluorescence micrographs (Figure 1B), which further emphasised the lower mannoprotein and chitin content of the NaCl-grown cells. The micrographs also showed that bright concanavalin A-bound FITC foci observed on the cell wall of cells grown in standard medium, likely corresponding to local mannoprotein clusters, were absent in NaCl-grown cells. Thus, growth in 1 M NaCl strongly affects the composition of the *D. hansenii* cell wall, decreasing its carbohydrate content and altering its surface organisation.

**Figure 1.**
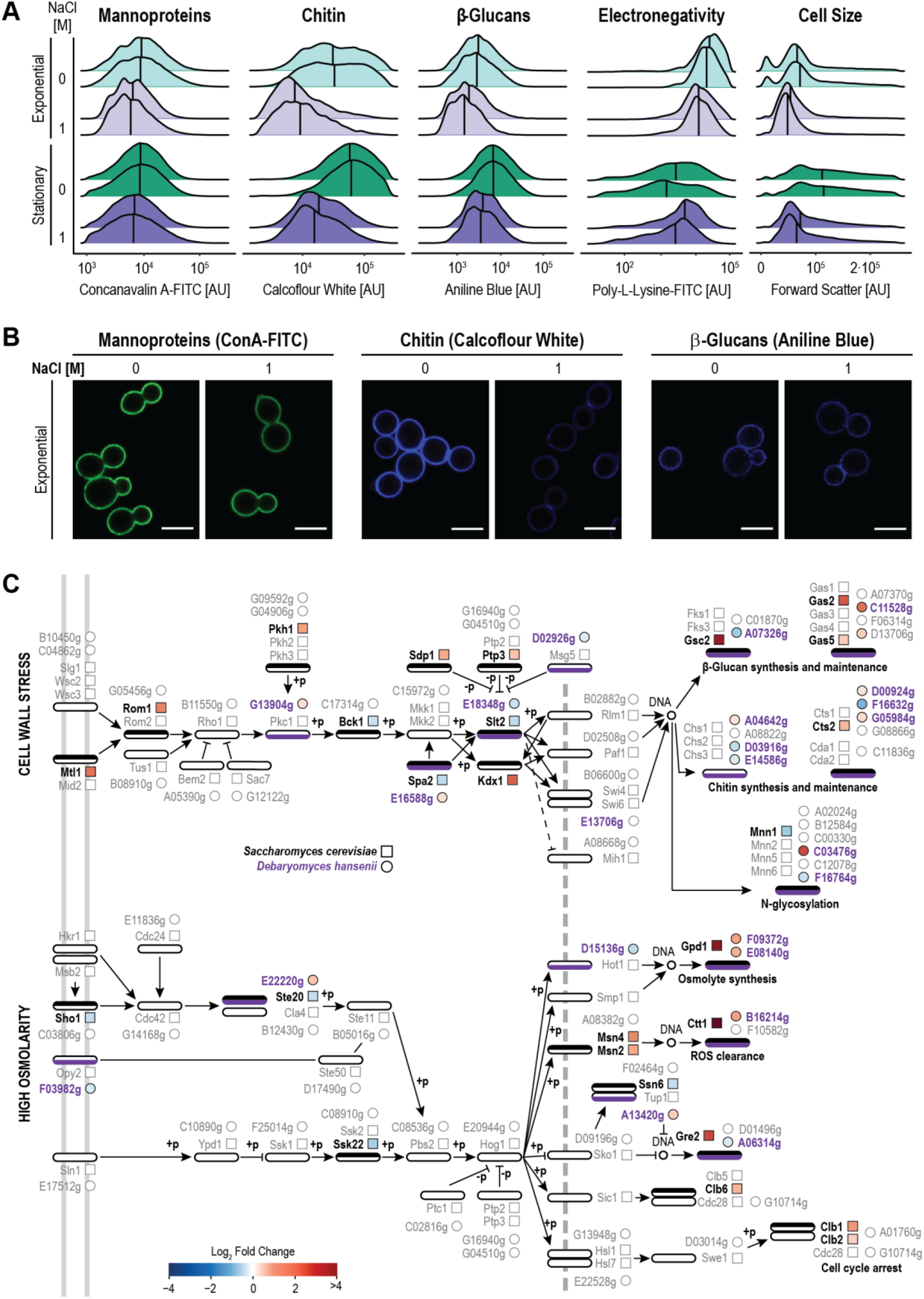
In a NaCl-supplemented medium, *D. hansenii* restructures its cell wall. (A) Flow-cytometry profiles of *D. hansenii* grown either in standard medium (0 M NaCl) or medium supplemented with 1 M NaCl, sampled in exponential (2 OD_600_/ml) and stationary phase. Cells were stained with fluorophores binding to mannoproteins (concanavalin A-bound FITC), chitin (Calcofluor White), and β-glucans (Aniline Blue). Electronegativity, i.e., cell-surface charge, was estimated with poly-L-lysine-bound FITC and cell size with forward scatter (FSC). Distributions show fluorescence intensity (arbitrary units, AU) for >10^4^ cells/sample. (B) Representative confocal micrographs of exponential phase cells grown in either standard medium or medium supplemented with 1 M NaCl, stained for mannoproteins (ConA-FITC), chitin (Calcofluor White), and β-glucans (Aniline Blue). Scale bars, 5 µm. (C) Schemes of the MAPK cascades (left) and downstream processes (right) detailing the response of *S. cerevisiae* (squares) and *D. hansenii* genes (circles) to cell wall stress and high osmolarity conditions (modified from KEGG pathway maps sce04011 and dha04011). Arrows denote activations, flat heads inhibitions, and ±p (de)phosphorylation. Next to each node, coloured markers report RNA-seq log_2_(fold-change) for each significantly differentially expressed orthologue (0.7 M vs 0 M NaCl for *S. cerevisiae*, 1.0 M vs 0 M NaCl for *D. hansenii*, adjusted p-value < 0.05). Non-differentially expressed genes are shown in grey. Bars above and below gene groups indicate if any gene from the group is differentially expressed (black for *S. cerevisiae* and purple for *D. hansenii* orthologues). The double grey line and hatched line depict the cell membrane and nuclear membrane, respectively.

To probe the molecular basis of the observed cell wall changes, we compared publicly available RNA-seq profiles of *D. hansenii* grown in a medium supplemented with 1.0 M NaCl versus the standard medium, and, in parallel, those of the model yeast *S. cerevisiae* grown in a medium supplemented with 0.7 M NaCl versus the standard medium. We focused on KEGG-derived mitogen-activated protein kinase (MAPK) signalling modules centred on plasma-membrane stress sensing, signal transduction, and transcriptional programmes that remodel the cell wall, maintain proteoglycan processing, produce osmolytes, detoxify ROS, and transiently restrain the cell cycle (Figure 1C).

Under salt stress, *S. cerevisiae* broadly induced the cell-wall integrity (CWI) pathway, with the transmembrane sensor Mtl1 (3.3-fold, p_adj_ = 5.3·10^-6^) and the protein kinase Kdx1 (4.0-fold, p_adj_ = 1.4·10^-20^) among the strongest responders. By contrast, *D. hansenii* showed little transcriptional induction of the corresponding pathway. Its most strongly induced component was the Pkc1 orthologue G13904g, which increased only 1.2-fold (p_adj_ = 0.02).

β-glucan remodelling was also affected differently. In *S. cerevisiae*, the induced CWI pathway significantly upregulated Gsc2 (1,3-β-D-glucan synthase catalytic subunit, 10.3-fold, p_adj_ = 9.5·10^-51^) as well as 1,3-β-glucanosyltransferases Gas2 (4.5-fold, p_adj_ = 1.0·10^-18^) and Gas5 (1.4-fold, p_adj_ = 0.03). By contrast, *D. hansenii* downregulated one of two 1,3-β-D-glucan synthase paralogues (A07326g, 1.8-fold, p_adj_ = 9.2·10^-9^), while upregulating two GAS paralogues (C11528g, 3.4-fold, p_adj_ = 4.2·10^-13^; D13706g, 1.4-fold, p_adj_ = 1.0·10^-3^). These expression changes are consistent with the observed decrease in *D. hansenii* β-glucan content, as reduced synthesis, coupled with enhanced processing, would thin the β-glucan network.

In *D. hansenii*, chitin homeostasis followed the same downregulation pattern. In *D. hansenii*, one paralogue of chitin synthase (CHS) was mildly upregulated (A04642g, 1.2-fold, p_adj_ = 0.03) while two were downregulated (D03916g, 1.5-fold, p_adj_ = 1.1·10^-4^; E14586g, 1.2-fold, p_adj_ = 4.3·10^-3^). Moreover, two *D. hansenii* chitinases were mildly upregulated (D00924g, 1.2-fold, p_adj_ = 0.03; G05984g, 1.3-fold, p_adj_ = 0.02), and one was downregulated (F16632g, 2.5-fold, p_adj_ = 2.8·10^-16^), indicating active rebalancing of chitin synthesis and turnover. By contrast, salt did not affect *S. cerevisiae* CHS, but it induced its chitinase Cts2 (1.7-fold, p_adj_ = 1.3·10^-3^). These observations account for the *D. hansenii* weak chitin signal in NaCl-supplemented medium (Figures 1A and 1B).

Finally, the loss of bright ConA-FITC foci on the cell wall of *D. hansenii* correlated with strong induction of a mannosyltransferase in NaCl-supplemented medium (C03476g, 5.2-fold, p_adj_ = 5.3·10^-7^), suggesting altered N- and O-mannan processing and surface distribution.

Together, this pathway-level comparison suggests a species-specific strategy. *S. cerevisiae* mounts a robust, multi-tier CWI response that promotes polysaccharide synthesis and remodelling, whereas *D. hansenii* shows limited transcriptional induction of the CWI pathway and selectively rewires β-glucan and chitin synthesis, as well as processing, thus shaping a thinner, reorganised cell wall, likely adapted for high-salinity growth.

We also examined the high-osmolarity glycerol (HOG) pathway, which is a related MAPK signalling module (Figure 1C). In both species, transcript levels of the core MAPK cascade were largely unchanged, consistent with its predominantly post-translational control (Blomberg, 2022). However, the downstream transcriptional programmes diverged. In *S. cerevisiae*, HOG activation drove a robust osmoadaptive response, strongly inducing glycerol synthesis via glycerol-3-phosphate dehydrogenase Gpd1 (11.3-fold, p_adj_ = 3.9·10^-55^), defence against reactive oxygen species via catalase Ctt1 (64.4-fold, p_adj_ = 4.9·10^-157^), and detoxification of reactive carbonyls via methylglyoxal reductase Gre2 (5.4-fold, p_adj_ = 8.8·10^-37^). In contrast, *D. hansenii* mounted a milder response. Its Gpd1 paralogues were only modestly upregulated (F09372g, 2.1-fold, p_adj_ = 1.8·10^-5^; E08140g, 1.8-fold, p_adj_ = 1.6·10^-6^). Of the two catalases, B16214g was mildly induced (2.1-fold, p_adj_ = 2.9·10^-5^) while F10582g remained unchanged (1.04-fold, p_adj_ = 0.79). Among the two Gre2 paralogues, A06314g was downregulated (1.24-fold, p_adj_ = 0.01), whereas D01496g was unaffected (1.03-fold, p_adj_ = 0.79). Moreover, NaCl altered the expression of cell-cycle-associated transcripts *CBL1-2-6* in *S. cerevisiae* but not of their homologues in *D. hansenii*. Together, these observations further underscore a species-specific osmoadaptive strategy: *S. cerevisiae* engages a robust HOG-driven programme that enhances glycerol synthesis, ROS quenching, and carbonyl detoxification, whereas *D. hansenii* mounts a comparatively weak downstream transcriptional response, with modest induction of osmolyte and stress-clearance genes, and minimal effects on the cell cycle.

### 3.2. Transcriptome-guided selection of *D. hansenii* promoters

Given the atypical remodelling of the *D. hansenii* cell wall in NaCl-supplemented medium, we wondered whether this yeast could serve as a favourable chassis for hypersaline surface display. Because molecular tools validated at high salt are scarce for *D. hansenii*, we first sought promoters that remain functional or become strongly induced under hypersaline conditions. We re-analysed a public RNA-seq dataset comparing *D. hansenii* cells grown in standard medium versus medium supplemented with 1 M NaCl. Of 6,425 expressed genes, 992 (15.4 %) were significantly upregulated, while 968 (15.1 %) were downregulated (|log_2_FC| > 0, p_adj_ < 0.10, Figure 2A).

**Figure 2.**
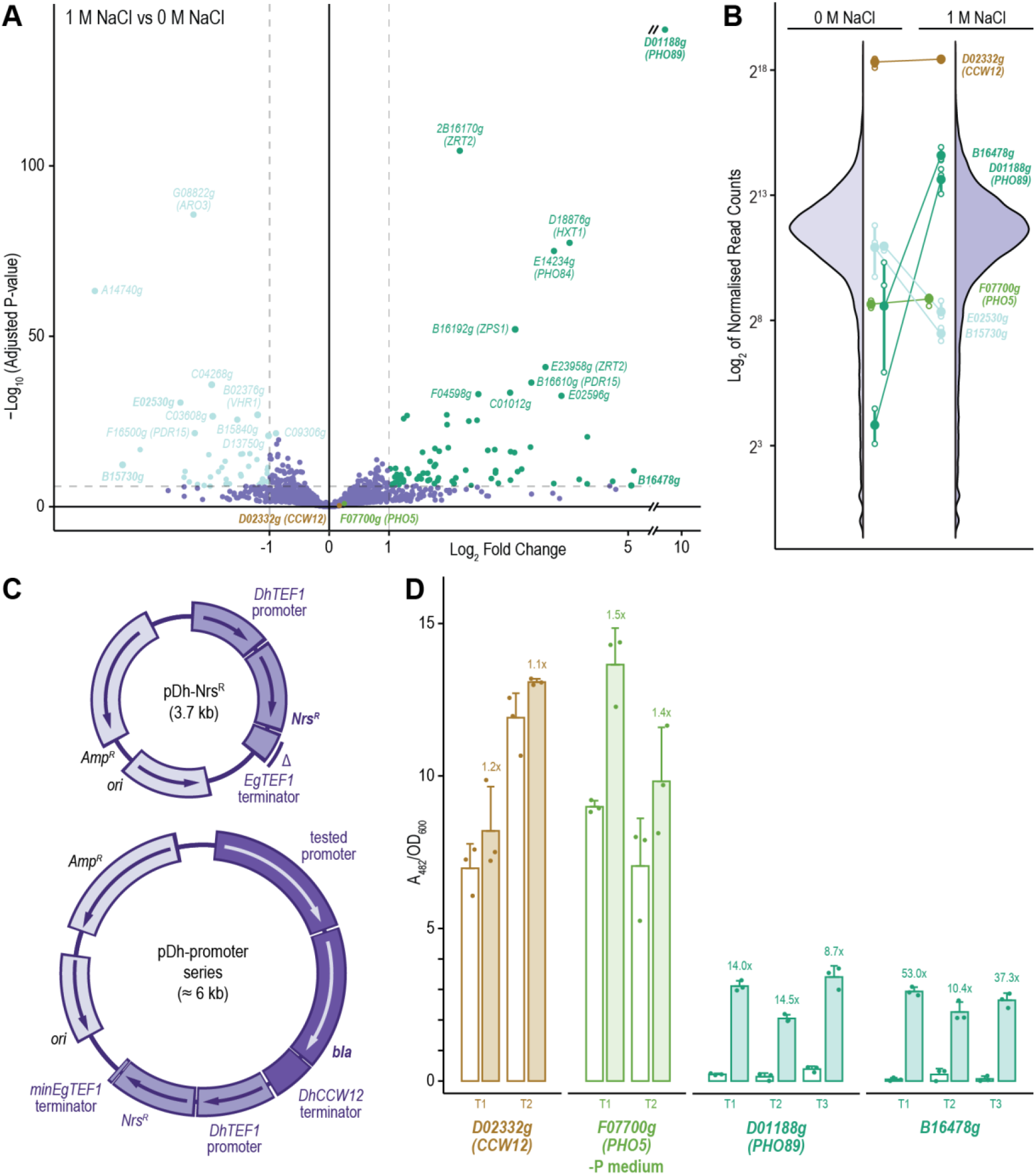
Developing strong constitutive and inducible *D. hansenii* expression vectors. (A) Volcano plot summarising differences in gene expression in 1 M NaCl-supplemented vs standard medium. The x-axis shows log_2_(fold change) (log_2_FC) and the y-axis shows −log_10_(adjusted P-values). Points are coloured by significance and expression direction (purple, not significant; dark green, upregulated; light green, downregulated). The strongest responders are annotated with their *D. hansenii* locus IDs and, when available, *S. cerevisiae* orthologue names in parentheses. Also highlighted are the *CCW12* orthologue *D02332g* (*DhCCW12*), which is the most strongly constitutively expressed gene in *D. hansenii*, and the *PHO5* orthologue *F07700g* (*DhPHO5*), which is strongly expressed in phosphate-free medium during the stationary phase. An x-axis break denotes extreme effect sizes. (B) Violin plots showing the genome-wide distribution of log_2_-normalised read counts for each gene in NaCl-supplemented and standard medium. Small and large circles denote biological replicates and mean values, respectively, and horizontal lines link the same gene across conditions. The two most up- and down-regulated genes (dark and light green, respectively) are highlighted alongside *DhCCW12* and *DhPHO5*. The colour scheme matches the one in panel A. (C) Maps of the initial transformation vector pDh-Nrs^R^ and plasmid series pDh-promoter used to evaluate promoter strength. *DhTEF1* – *D. hansenii TEF1* promoter, *Nrs^R^*– nourseothricin-resistance cassette, codon-optimised for expression in *D. hansenii*, *EgTEF* terminator – *TEF* terminator from *Eremothecium gossypii*, *Amp^R^* – *E. coli*-expressed ampicillin-resistance marker, *ori* – *E. coli* plasmid origin of replication, *bla* – yeast-codon-optimised β-lactamase used as the reporter gene for testing promoter activity, *DhCCW12* terminator – terminator of the *D. hansenii* gene *CCW12*, *minEgTEF1* terminator – minimal *EgTEF1* terminator variant engineered to remove cryptic *D. hansenii* replication origin (Δ). (D) Promoter activities quantified with a yeast-codon-optimised β-lactamase reporter. Bars show mean enzymatic activity for each promoter in standard medium (open bars) and in medium supplemented with 1 M NaCl (shaded). Points denote individual biological replicates, error bars indicate standard deviations, and numbers compare expression level relative to cells grown in standard medium. Colours identify promoters as in panel A.

Guided by these results, we focused on several promoters. Among the induced transcripts, the most responsive was the *D. hansenii* orthologue of *S. cerevisiae PHO89* (*D01188g*), which in *S. cerevisiae* encodes a plasma membrane Na^+^/Pi cotransporter (861-fold increase, p_adj_ = 1.0·10^-140^). We therefore chose the *DhPHO89* promoter (*pDhPHO89*) as a candidate NaCl-inducible element, together with the promoter of another salt-induced gene, *B16478g* (37.3-fold, p_adj_ = 8.0·10^-6^). As a robust constitutive promoter, we selected the most highly expressed gene in both media, *D02332g* (*pDhCCW12*, Figure 2B), one of three putative *D. hansenii CCW12* paralogues. Finally, we also tested the promoter of the *D. hansenii PHO5* orthologue (*pDhPHO5*), whose *S. cerevisiae* homologue is strongly induced in phosphate-free medium during the stationary phase.

### 3.3. *EgTEFt* confers cryptic replication activity in *D. hansenii*

To test the functionality and strength of these promoters, we constructed a transformation vector pDh-Nrs^R^ (Figure 2C). We codon-optimised for *D. hansenii* and de novo synthetised an open reading frame (ORF) encoding nourseothricin acetyltransferase, conferring resistance to nourseothricin (clonNAT). We inserted the ORF into a pFA6a plasmid backbone under the control of the strong *D. hansenii TEF1* promoter (*pDhTEF1*) while retaining the terminator of the *Eremothecium gossypii TEF* gene (*EgTEFt*), as this terminator is standardly used in the yeast MX-marker series. After sequencing, we introduced the plasmid into *D. hansenii* by electroporation.

Although pDh-Nrs^R^ was designed as an integrative vector, in *D. hansenii* it yielded unexpectedly high transformation efficiency. We consistently obtained several thousand small transformant colonies per plate that failed to grow in liquid medium supplemented with clonNAT, even when the plasmid was introduced in circular form (2.6 ± 1.1·10^6^ μg^-1^ DNA). Similarly high transformation efficiencies were observed following transformation with pDh-Nrs^R^ linearised by PvuI (2.1 ± 1.2·10^6^ μg^-1^ DNA) or with PCR-amplified DNA generated using non-phosphorylated primers (5.5 ± 1.7·10^6^ μg^-1^ DNA). These results suggest that *D. hansenii* efficiently circularises exogenous DNA, even when the process requires more than a simple ligation.

We tried to avoid the small-transformant phenotype by reducing the marker dosage. We lowered Nrs^R^ expression by deleting 430 bp of *pDhTEF1* to generate a minimal 100-bp promoter (*pminDhTEF1*) containing only the TATA box, without upstream activating sequences. However, this reduction in promoter strength did not measurably affect transformation efficiency nor change colony size or growth.

The small-transformant phenotype was reminiscent of abortive transformants described in *S. cerevisiae*, in which incoming DNA is stabilised and transcriptionally active but unable to replicate. Alternatively, pDh-Nrs^R^ could harbour a *D. hansenii*-compatible cryptic replication origin, because of which the plasmid would be inefficiently replicated and/or segregated between the mother and daughter cells. To distinguish between the two cases, we performed a series of targeted deletions that identified a 180-bp region required for high transformation efficiency, corresponding to part of *EgTEFt*. Deletion of this fragment markedly reduced transformation efficiency, with transformants obtained only when the plasmid was linearised with PvuI (6.2 ± 1.6·10^2^ μg^-1^) or introduced as a PCR amplicon (6.7 ± 2.8·10^3^ μg^-1^), but not when used in circular form. These results strongly suggest that *EgTEFt* provides origin-like activity enabling limited autonomous replication in *D. hansenii*.

### 3.4. Characterising *D. hansenii* promoters using a β-lactamase reporter

Next, we constructed a series of integrative reporter plasmids derived from pDh-Nrs^R^, with truncated *EgTEFt*, to quantify promoter strength and inducibility (the pDh-promoter series, Figure 2C bottom). We initially aimed to use a classical *PHO5*-based *p*-nitrophenyl phosphate reporter system widely applied in *S. cerevisiae*. However, our preliminary experiments revealed substantial endogenous phosphatase activity in *D. hansenii* even in a phosphate-rich medium, rendering such measurements unreliable. We therefore turned to an intracellular β-lactamase-based reporter, whose expression we evaluated on digitonin-permeabilised cells using a nitrocefin assay. We cloned each promoter of interest upstream of the β-lactamase ORF, PCR-amplified the resulting expression cassette together with the Nrs^R^ selection marker and transformed *D. hansenii* with the amplicon. For each promoter, we analysed at least two independent transformants to assess positional effects arising from random chromosomal integration (Figure 2D).

Consistent with transcriptomic data, *pDhCCW12* (*pD02332g*) drove strong, salt-insensitive expression in the exponential phase (6.8 vs 8.0 A_482_/OD_600_ in control vs 1 M NaCl, respectively; p = 0.29). Notably, expression levels differed substantially between integration events, with a 72 % difference observed between two independent transformants (6.8 ± 0.8 vs 11.7 ± 1.0 A_482_/OD_600_, p = 3.1·10^-3^), indicating a strong influence of genomic context on promoter output. The *pDhPHO5* (*pF07700g*) promoter drove high stationary-phase expression in phosphate-free medium, with increased activity in NaCl-supplemented medium (8.7 ± 0.3 vs 13.3 ± 1.2 A_482_/OD_600_, p = 0.06).

The assay further confirmed that *pDhPHO89* (*pD01188g*) and *pB16478g* are NaCl-inducible promoters, albeit weaker than *pDhCCW12* and *pDhPHO5*. Both promoters showed minimal activity in standard medium (0.1-0.4 and 0.1-0.2 A_482_/OD_600_, respectively), but were significantly induced in NaCl-supplemented medium, reaching 2.1-3.4 and 2.3-2.9 A_482_/OD_600_ (p = 0.04-3.2·10^-3^ and 0.2-2.0·10^-3^, respectively). Together, these data establish a compact promoter toolkit for *D. hansenii* comprising a strong salt-insensitive promoter (*pDhCCW12*), a strong stationary-phase promoter (*pDhPHO5*), and two NaCl-inducible promoters (*pDhPHO89* and *pB16478g*), suitable for controlled gene expression under hypersaline conditions.

### 3.5. In silico characterisation of *D. hansenii* Ccw12 paralogues

To implement functional surface display in *D. hansenii*, we needed to identify efficient signal peptide and anchoring sequences. We focused on *DhCCW12*, the most highly expressed gene in *D. hansenii*, whose homologue in *S. cerevisiae* encodes a highly abundant cell wall protein that is covalently anchored to the cell wall via a C-terminal GPI anchor. The two proteins share 36 % amino acid identity. We reasoned that surface display tags derived from such a highly abundant protein would be evolutionarily optimised for efficient secretion and covalent cell-surface anchoring. Interestingly, *DhCCW12* is one of three putative *CCW12* paralogues in *D. hansenii*, with the remaining two (*A03498g* and *G06182g*) expressed at approximately 5 % of the *DhCCW12* level (Figure 3A).

**Figure 3.**
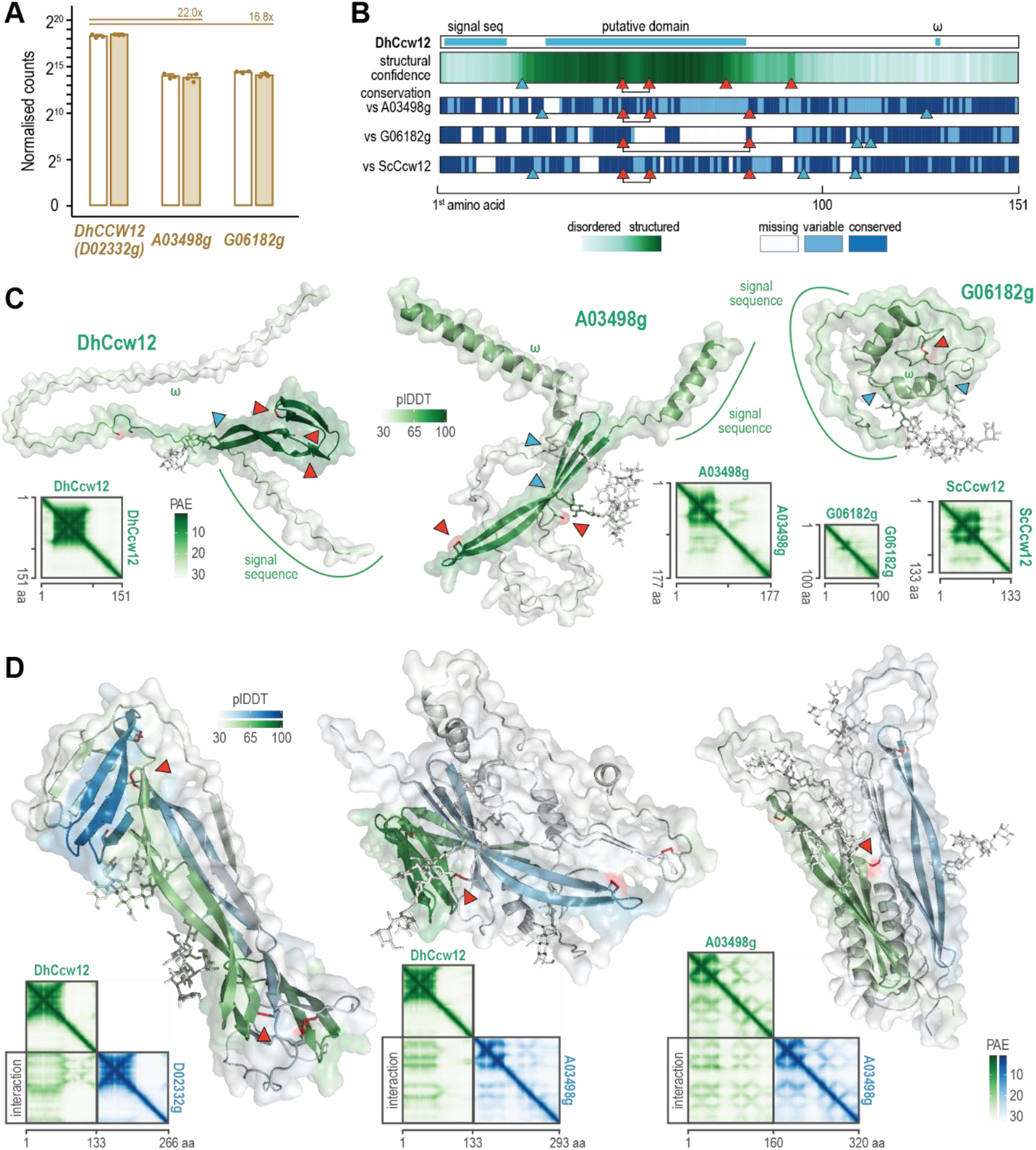
Sequence features and structural conservation of *D. hansenii* Ccw12 paralogues. (A) Normalised RNA-seq read counts for the three *D. hansenii CCW12* paralogues (*DhCCW12/D02332g*, *A03498g*, and *G06182g*) in standard (open bars) and NaCl-supplemented medium (shaded). Bars represent mean expression levels across biological replicates, with individual data points overlaid. Numbers indicate fold differences relative to *DhCCW12*. (B) Schematic overview of predicted sequence features and conservation across Ccw12 paralogues. Tracks show the signal peptide, putative structured domain, and the predicted GPI-anchor ω-site (ω) for DhCcw12, as well as sequence conservation relative to A03498g, G06182g, and *S. cerevisiae* Ccw12. Structural confidence predicted by AlphaFold3 in the form of local confidence scores per amino acid residue (pLDDT) is shown along the sequence, denoting predicted disordered regions (light green) versus structured regions (dark green). Blue triangles point to predicted N-glycosylation sites and red triangles to conserved cysteine residues. Predicted intrastrand disulfide bonds are shown as black lines between cysteine residues. (C) AlphaFold3-predicted monomeric structures of DhCcw12, A03498g, and G06182g, coloured by pLDDT scores, with marked signal peptides, GPI-anchor ω-residues, potential N-glycosylation sites and conserved cysteines. Insets show predicted aligned error (PAE) plots for each protein, illustrating global interaction confidence, with light and dark green denoting low and high confidence, respectively. (D) AlphaFold3-predicted homo- and heterodimeric assemblies of DhCcw12 and A03498g, highlighting domain-domain interactions and predicted intermolecular disulfide bonds. Structures are coloured by pLDDT, and corresponding PAE plots illustrate confidence in interchain contacts.

To identify relevant sequence features, we predicted signal peptides, GPI-anchor sites, putative N-glycosylation motifs, and three-dimensional structures for all three paralogues (Figure 3B). All paralogues contained a clearly predicted signal peptide, a conserved GPI transamidase-targeted ω-site, at least one N-glycosylation site, and at least two of the three cysteine residues conserved in ScCcw12. AlphaFold3 further predicted that DhCcw12 and A03498g, like ScCcw12, contain a structured domain stabilised by an intrachain disulfide bond (Figure 3C). In contrast, the 50 aa shorter paralogue G06182g was predicted as completely disordered, lacking one of the cysteine residues required for disulfide bond formation. Thus, DhCcw12 (D02332g) and A03498g recapitulate the key structural features of ScCcw12.

We previously hypothesised that ScCcw12 functions as a glycan-rich adaptor protein that uses its structured interaction domain to form homo- and heterodimers (Martinić Cezar et al., 2024). We investigated whether similar behaviour could be expected for DhCcw12 and A03498g. Consistent with this model, AlphaFold3 predicted that both proteins could form homo- and heterodimers via domain-domain interactions (Figure 3D). Such assemblies were also predicted to be stabilised by one or two intermolecular disulfide bonds. Collectively, these findings indicate that DhCcw12 and A03498g recapitulate all salient structural and oligomerisation features of ScCcw12.

### 3.6. DhCcw12 enables functional surface display in *D. hansenii*

We next constructed a series of pDh-reporter plasmids for surface display of reporter enzymes using either C- or N-terminal anchoring strategies (Figure 4A). In all constructs, the surface-display cassette was placed under the control of the robust *DhPHO5* promoter and the *DhCCW12* terminator, and included a triple HA epitope tag for immunoblot detection. For C-terminal anchoring, we inserted *E. coli* β-lactamase between the previously identified DhCcw12 signal peptide and C-terminal GPI-anchoring sequence (Figure 4B). Following processing, this strategy increased the apparent molecular weight of the reporter enzyme by <2 kDa (Figure 4C). To generalise our findings across enzyme classes, we also tested two additional reporter enzymes for which cell-impermeable chromogenic substrates are available and inserted them into the reporter cassettes, namely acetylxylan esterase (AXE) from *Bacillus pumilus* and lipase A from *Bacillus subtilis*. For N-terminal anchoring, we employed a previously optimised β-lactamase fusion protein based on *S. cerevisiae* Pir2 (Hsp150), selected as the best-performing among 12 tested variants (Martinić Cezar et al., 2023).

**Figure 4.**
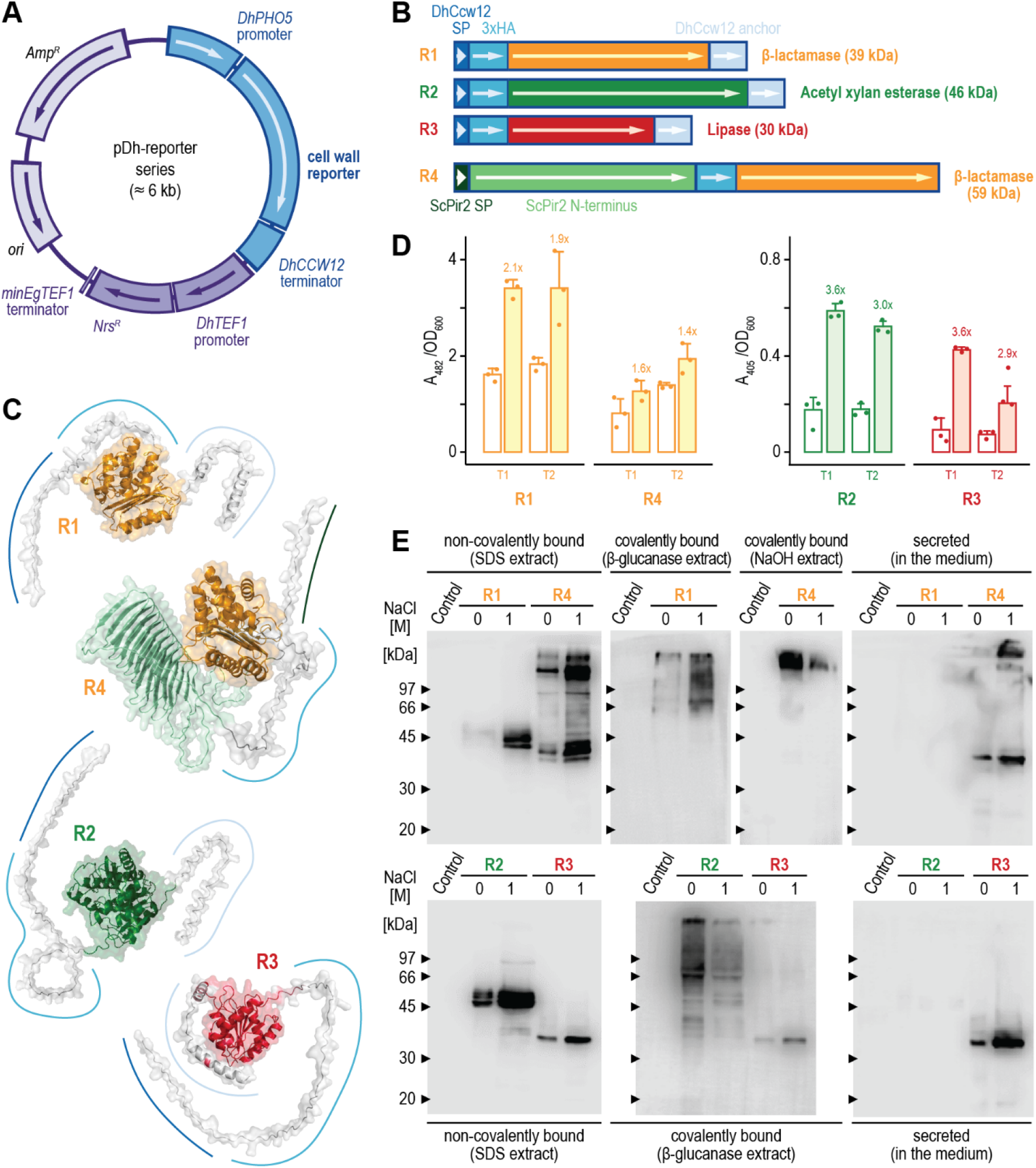
Implementing *D. hansenii* surface display. (A) Schematic map of the pDh-reporter plasmid series designed to express *D. hansenii* reporters for surface display. The reporter cassette is driven by the *D. hansenii PHO5* promoter (*DhPHO5*), terminated by the *DhCCW12* terminator, and includes a triple HA epitope tag for immunodetection. The vector backbone contains the *Nrs^R^* selection cassette under the control of the *DhTEF1* promoter, a minimal *EgTEF1* terminator that lacks replication activity in *D. hansenii*, and bacterial *Amp^R^*and *ori* elements for replication in *E. coli*. (B) Design of surface-display reporters. Reporters R1-R3 carry the DhCcw12 signal peptide (SP) and are C-anchored through GPI attachment, while R4 is N-anchored via an ScPir2-based fusion. Reporter enzymes include β-lactamase, acetylxylan esterase (AXE), and lipase A. Theoretical molecular masses of the processed reporters are indicated in parentheses. (C) AlphaFold3-predicted structures of surface-display reporters R1-R4, with the reporter domains highlighted. Colours match ORF segments in panel B. (D) Enzymatic activity of surface-displayed reporters measured by cell-impermeable chromogenic substrates nitrocefin (β-lactamase) and *p*-nitrophenyl acetate (AXE and lipase A). Activities are shown for cells grown in standard phosphate-free medium (open bars) and in phosphate-free medium supplemented with 1 M NaCl (shaded bars). Bars represent mean activity (A_482_ or A_405_) normalised to cell density (OD_600_), points indicate individual biological replicates, error bars denote standard deviations, and numbers compare activities relative to those of cells grown in standard medium. (E) Anti-HA immunoblot of cell wall protein fractions. Cell wall proteins were isolated as noncovalently bound (SDS extract) and covalently bound (β-glucanase or NaOH extracts) protein fractions, as well as a protein fraction from culture supernatant. The fractions were analysed by SDS-PAGE and probed for cell wall reporters. Samples were derived from cells grown in standard medium (0 M NaCl) or NaCl-supplemented medium (1 M). Molecular masses of protein standards are denoted on the left.

Following expression of the surface-display cassettes, we quantified activities of the displayed reporters (Figure 4D). All reporters were functional and consistently showed higher activity in cells grown in NaCl-supplemented medium, with an increase of >80 % for β-lactamase (3.41 vs 1.83 A_482_/OD_600_, p = 0.06) and >200 % for acetylxylan esterase (AXE; 0.51 vs 0.17 A_405_/OD_600_, p = 3·10^-4^) and lipase (0.42 vs 0.12 A_405_/OD_600_, p = 0.02). Moreover, C-terminal anchoring via DhCcw12 outperformed N-terminal anchoring based on ScPir2, particularly under NaCl-supplemented conditions (3.41 vs 1.94 A_482_/OD_600_, p = 0.05).

To determine which cell wall fraction the reporter enzymes localised to, we performed an anti-HA immunoblot (Figure 4E). All reporters were detected in both covalently and noncovalently bound protein fractions, with C-anchored lipase and N-anchored β-lactamase additionally detected in the culture medium. In line with activity assays, stronger signals were consistently observed in cell wall fractions derived from cells grown in NaCl-supplemented medium. Moreover, covalently bound reporters typically appeared as multiple bands with higher apparent molecular masses, likely reflecting heterogeneity in post-translational processing. In contrast, noncovalently bound fractions migrated predominantly as single bands or closely spaced doublets near their predicted molecular weights, consistent with little or no glycosylation.

## 4. Discussion

In this work, we engineered the halotolerant yeast *D. hansenii* into a host for functional surface display under hypersaline conditions. We showed that growth in 1 M NaCl-supplemented medium remodels the cell wall and increases the activity of three unrelated surface-displayed enzymes. To build this system, we identified native promoters for constitutive, stationary-phase, and NaCl-inducible expression, characterised an unexpected cryptic replication element, and adapted the highly expressed cell wall protein DhCcw12 as a secretion and GPI-anchoring module. Together, these findings show that the adaptations allowing *D. hansenii* to thrive at high salinity also support cell-surface biocatalysis under conditions poorly tolerated by conventional production hosts.

Compared to cells grown in standard medium, *D. hansenii* cells grown in 1 M NaCl-supplemented medium build a thinner cell wall. This result is counterintuitive, as a salinity-facing yeast would be expected to reinforce its cell wall, as several halophilic fungi do, e.g., the basidiomycete *Wallemia ichthyophaga* (Kralj Kunčič et al., 2010), the black yeasts of the genus *Trimmatostroma* (Kogej et al., 2006), and the marine fungus *Dendryphiella salina* (Clipson et al., 1989). Yet the cell wall thickening is not the universal response, with the model yeast *S. cerevisiae* remodelling rather than thickening its cell wall under chronic salt exposure (Rodríguez-Peña et al., 2010), because the structure’s stiffness depends more on β-glucan-chitin cross-linking than on the abundance of any single polysaccharide (Dague et al., 2010; Ribeiro et al., 2021). On the other hand, halotolerant fungi favour the cell wall’s flexibility over its reinforcement. For example, in NaCl-supplemented medium, the moderately halotolerant fungi *Aspergillus flavus* and *Penicillium roquefortii* (Abu-Seidah, 2007), the black yeast *Aureobasidium pullulans* (Kralj Kunčič et al., 2010) and the halotolerant yeast *Zygosaccharomyces rouxii* (Wang et al., 2020) thin their cell wall, with *Z. rouxii* also lowering its mannan content (Dakal et al., 2014; Hamada et al., 1984; Hosono, 1992). For these species, a more pliable cell wall likely helps NaCl tolerance, as *Z. rouxii* isolate with a rigid cell wall tolerates salt worse than its counterpart with an elastic cell wall (Přibylová et al., 2007). Following its halotolerant phenome, *D. hansenii* responds similarly.

However, from a broader perspective, the *D. hansenii* cell wall thickness depends on the precise NaCl concentration in the medium. Electron micrographs show that the *D. hansenii* cell wall thins at 0.6 M NaCl but thickens at 2.4 M NaCl (Ochoa-Gutiérrez et al., 2022), suggesting that the actual response depends on whether the NaCl concentration falls within the range to which the species is well adapted. Thus, the observed high-salinity contrast, with some fungi thickening and others thinning their cell wall, likely reflects a trade-off between cell wall rigidity and flexibility, determined by how stiffness is built into the cell wall and by the severity of the salt stress.

Under sustained 1 M NaCl, *D. hansenii* activates stress response more modestly than *S. cerevisiae*. Specifically, the core high-osmolarity cascade, being post-translationally controlled, remains transcriptionally near-silent in both species. Instead, the species diverge downstream, with *S. cerevisiae* strongly driving glycerol synthesis, mounting ROS defence, and detoxifying carbonyls, whereas *D. hansenii* responds more mildly. For example, although *DhHOG1* and *DhMSN2* transcripts can respond to saline or oxidative stress (Ramos-Moreno et al., 2019), neither shifted under the sustained 1 M salinity we examined. This downstream split complements earlier reports that activated DhHog1 remains phosphorylated and active longer than ScHog1 (Sánchez et al., 2020), but enters the nucleus more slowly and lingers in the cytoplasm for much longer during osmostress (Sharma et al., 2005). *D. hansenii* therefore appears to reprogram gene expression sparingly, narrowing DhHog1-driven transcription, keeping the kinase active in the cytoplasm, and leaning instead on constitutive, post-translational, and partly Hog1-independent cytoplasmic routes (de la Fuente-Colmenares et al., 2024).

Such gene-expression restraint may begin even earlier, at the level of osmosensing. Unlike *S. cerevisiae*, *D. hansenii* requires an intact cell wall to fully mount its halotolerance response (Thomé, 2007), suggesting that it senses osmolarity through ligand-receptor contact between cell-wall components and a plasma-membrane receptor, rather than through the membrane alone. This cell-wall-dependent architecture could reinforce the above-described transcriptional dampening. Specifically, as *D. hansenii* remodels its cell wall under high salinity, the remodelling might alter the contact between the cell wall and the sensor, lowering sensor output and further restricting downstream programmes. Such a reduction carries little risk for *D. hansenii*, for which salinity is a native condition and in which NaCl imposes osmotic rather than ionic stress, sodium accumulates to high intracellular levels and can substitute for potassium (Prista et al., 1997), and salt even protects the cell against further stresses (Michán et al., 2013; Navarrete et al., 2009). Thus, rather than reduced sensitivity, the proposed sensor damping likely reflects homeostatic adaptation to high salinity.

*D. hansenii* is an appealing production host, but its genetic toolkit lags behind. Until recently, work on its promoters was confined to a limited set of heterologous expression tests (Maggi and Govind, 2004). A modular collection has since begun to close that gap, supplying genomic integration loci and constitutive promoters that span three orders of magnitude in expression, all derived by homology to *S. cerevisiae* promoters (Li et al., 2025). Inducible promoters, though, remain scarce, and homology to *S. cerevisiae* gives little indication of which *D. hansenii* promoters respond to a given condition. We therefore mined *D. hansenii*’s own transcriptome instead, re-analysing the DebaryOmics NaCl dataset (Navarrete et al., 2022b), reproducing the reported salt-responsive signature, and identifying NaCl-inducible promoters alongside a strong constitutive one.

*DhPHO89*, the most strongly NaCl-induced gene we found, is the orthologue of a phosphate-starvation gene, yet our medium was phosphate-replete. In *S. cerevisiae*, *PHO89* encodes the Na^+^-coupled high-affinity phosphate transporter, the alkaline-pH counterpart of the H^+^-coupled Pho84, induced by phosphate limitation through Pho4 (Serra-Cardona et al., 2014). However, *S. cerevisiae PHO89* is also induced by the calcineurin-Crz1 pathway, which responds to saline and alkaline stress independently of phosphate, through the regulatory motifs that also induce the transcription of *ENA1*, a Na^+^-extrusion ATPase (Serra-Cardona et al., 2014; Serrano et al., 2002). *DhPHO84* is also likely induced by the calcineurin-Crz1 pathway to import additional phosphate to regenerate ATP, run ion pumps, and synthesise glycerol, thus coping with increased salinity. Concurrently, as the Na^+^-coupled transporter uptakes phosphate via the Na^+^ gradient, DhEna1 extrudes the imported Na^+^ ions. Thus, under salt stress, *D. hansenii* likely uses sodium energetics to meet its increased phosphate needs.

In our attempts to expand *D. hansenii*’s genetic toolbox, we noticed that PvuI-linearised and PCR-amplified plasmid backbone transformed *D. hansenii* as efficiently as did the intact circular plasmid, suggesting highly efficient end joining, i.e., circularisation of linear DNA. The circularisation of transforming DNA has also been previously observed in *S. cerevisiae*, where it underlies abortive transformation (Yap and Schiestl, 1995), and in *Candida glabrata*, where it proceeds via non-homologous end joining (Corrigan et al., 2013). However, *S. cerevisiae* prefers to circularise cohesive and phosphorylated ends, with the frequency of abortive transformation reflecting how readily the termini religate, whereas in *C. glabrata* blunt self-closure occurs but is rare. On the other hand, *D. hansenii* showed no such dependence, suggesting that its DNA-repair machinery efficiently converts both blunt and 5′-hydroxyl ends into ligatable junctions.

We also determined that *EgTEFt*, the terminator of the *E. gossypii TEF* gene, in *D. hansenii* functions as an inefficient, cryptic replication origin. This 200 bp AT-rich (33 % GC) terminator fits the general mould of the budding yeast autonomous replication sequence (ARS). For example, Fukuhara (2006) identified 80-bp AT sequences that support replication in *S. cerevisiae*, *K. lactis* or *T. delbrueckii*, and lack a shared motif or *S. cerevisiae* ARS consensus sequence (ACS), which suggests their ARS activity stems from their AT-rich nature. *D. hansenii* ARSs fit this pattern, as they are likewise extended AT-rich elements that carry no ACS (Minhas et al., 2009). Relatedly, Fukuhara’s 80-bp-carrying plasmids produced colonies of different sizes, the smallest of which mirror our weakly maintained *EgTEFt* plasmids. Thus, our results show that a transcriptional terminator can double as a cryptic replication origin, turning an intended integrative marker into an unstable, background-generating episome.

To construct novel expression vectors, we relied on native *D. hansenii* promoters identified from DebaryOmics RNA-seq data (Navarrete et al., 2022b) and tested them across growth phases and salt conditions. Previous studies either tested the expression of mostly heterologous *S. cerevisiae* promoters carried by unstable ARSD-based replicative plasmids (Maggi and Govind, 2004) or chose them based on homology to *S. cerevisiae* housekeeping genes, integrated them at defined loci, and characterised them in the late-exponential phase (Li et al., 2025). Our complementary approach focused on the salt-responsive and stationary-phase behaviour that homology to *S. cerevisiae* genes cannot predict.

We developed two strong expression vectors, based on promoters of *DhCCW12* and *DhPHO5*, designed for expression in the exponential and stationary phase, respectively. While promoter strength is typically measured in exponential cells, surface display targets high cell density, so expression in the stationary phase is critical for prolonged surface display. Stationary-phase promoters address this need, e.g., the promoter of the stationary-phase-expressed Sed1 for *S. cerevisiae* surface display (Inokuma et al., 2014). Yet, stationary-phase expression cannot be reliably inferred from exponential or *S. cerevisiae* data, as *D. hansenii* is a Crabtree-negative yeast lacking diauxic glycolytic repression (Sánchez et al., 2006). *DhPHO5* is convenient because it is highly active in the stationary phase in a medium lacking phosphate and because it directs the expression of a phosphatase targeted to the periplasm and cell wall, making it an appropriate driver for cell wall-anchored cargo.

As our constructs are integrated at random chromosomal loci, the measured promoter strength is heavily influenced by genomic context. This phenomenon is well known, having been noted in *D. hansenii* (Li et al., 2025) and systematically surveyed in *S. cerevisiae*, with studies showing that promoter strength can vary 8.7-fold (Bai Flagfeldt et al., 2009) or even up to 13-fold, with the promoter, reporter, or carbon source affecting expression less than the position of the expression cassette (Wu et al., 2017). This variation is influenced by local chromatin and the density of essential genes (Gui et al., 2021). Having analysed only two integrants per promoter, we observe a more modest effect, but one that also affects the inducibility fold change. However, the characterised NaCl-inducible promoters offer a starting point for further engineering. Stress-responsive elements have been assembled into strong, condition-inducible synthetic promoters in *S. cerevisiae* (Rajkumar et al., 2016), and the same route could strengthen the response of the here-reported promoters.

We developed a C-terminal anchoring surface display based on the most highly expressed *D. hansenii* gene, which encodes the cell wall protein DhCcw12, one of three ScCcw12 homologues in *D. hansenii*. We selected it because ScCcw12 is an abundant GPI-anchored mannoprotein that stabilises the cell wall at sites of active synthesis, and its loss leaves the cell wall disordered, over-chitinised, and lysis-prone (Ragni et al., 2011; Ragni et al., 2007). We previously proposed that ScCcw12 functions as a glycan-rich adaptor, folding into a small structured domain that enables homo- and heterodimerisation (Martinić Cezar et al., 2024). AlphaFold3 extends this model to DhCcw12, which shares only 36 % identity with ScCcw12 across more than 200 million years of divergence, predicting that DhCcw12 and A03498g dimerise through the same domain via intermolecular disulfide bonds, suggesting such an architecture is conserved and selected for. We also uncovered three *D. hansenii* homologues, a finding aligned with Saccharomycotina evolutionary trends, as yeast GPI-anchoring apparatus is conserved, but the cell wall proteins it delivers are often gained, lost, or multiplied in a clade-specific manner (Lozančić et al., 2021). Building on these insights, we developed the DhCcw12-based system, modelled on our *S. cerevisiae* minimal Ccw12-tag (Martinić Cezar et al., 2024), which performed well and could potentially be improved further in a *Dhccw12Δ* or similar background, as removing an abundant native cell wall protein can increase surface display capacity by freeing anchoring sites (Bamba et al., 2018) or thickening the cell wall (Inokuma et al., 2021).

We displayed three unrelated enzymes on the *D. hansenii* cell wall, which, to our knowledge, is the first surface display in this yeast. These results establish *D. hansenii* as a host for functional surface display using both native GPI-dependent C-terminal anchoring and heterologous Pir-based N-terminal anchoring (Martinić Cezar et al., 2023). The functionality of all three reporters also indicates that the DhCcw12 signal peptide and GPI-attachment sequence can accommodate proteins with different structures and catalytic mechanisms, suggesting that this anchor may have broad applicability.

Finally, our results also reveal NaCl supplementation as a simple means of substantially increasing surface-displayed reporter activity. As a halotolerant yeast, *D. hansenii* responds to salinity through extensive physiological adaptation involving its plasma membrane, secretory functions and cell-envelope-associated proteins (Navarrete et al., 2022b; Turk et al., 2007). Combined with stronger expression from the *DhPHO5* promoter in NaCl-supplemented medium, such adaptation could enhance recombinant protein folding, trafficking, stability, or cell-surface retention. Salt-dependent changes in cell wall architecture or porosity may also improve substrate access to displayed enzymes. Relatedly, immunoblotting detected the reporters in both covalently and noncovalently bound cell wall fractions, as well as in the culture medium, consistent with incomplete wall incorporation and the release of a proportion of the expressed protein (Kipnis et al., 2004). Together, these results support the use of *D. hansenii* in whole-cell biocatalysis, particularly in non-sterile, saline, or low-water-activity processes where its intrinsic robustness may offer an advantage over conventional yeast hosts (Andreu and del Olmo, 2020; Estrada et al., 2023b).

## 5. Conclusion

In this work, we characterised how *D. hansenii* remodels its cell wall in a medium supplemented with 1 M NaCl and used these insights to build a toolbox for C- and N-terminally anchored surface display. Salt-responsive promoters, together with the DhCcw12 signal peptide and GPI-anchoring sequence, supported the functional display of diverse reporter enzymes, with NaCl supplementation increasing their activities by >80 %, despite reducing cell-wall polysaccharide content. This combination of salt-compatible expression and cell wall anchoring establishes *D. hansenii* as a tunable, halostable surface display platform for whole-cell biocatalysis in saline or low-water-activity bioprocesses.

## Supplementary Material

Sequences of plasmids used in this study.

## Availability of data and materials

The datasets used and/or analysed during the current study are available from the corresponding author on reasonable request.

## Declaration of generative AI and AI-assisted technologies in the manuscript preparation process

During the preparation of this work, the authors used Claude Opus 4.8 in order to enhance readability and language. After using this tool, the authors reviewed and edited the content as needed and take full responsibility for the content of the published article.

## Authors’ contributions

AP: Methodology, Validation, Formal analysis, Investigation. TMC: Formal analysis, Investigation. BV: Resources, Writing - Review & Editing. RT: Resources, Supervision, Writing - Review & Editing. BŽ: Conceptualization, Methodology, Software, Formal analysis, Investigation, Writing - Original Draft, Writing - Review & Editing, Visualization, Supervision, Project administration, Funding acquisition. All the authors reviewed the manuscript.

## Acknowledgements

We thank David Gosset for his help with fluorescent microscopy and flow cytometry on the MO2VING platform (Centre de Biophysique Moléculaire, CNRS, Orleans).

## Literature

Abu-Seidah, A.A., 2007. Effect of Salt Stress on Amino Acids, Organic Acids and Ultrastructure of Aspergillus flavus and Penicillium roquefortii. Int. J. Agric. & Biol. 9(3), 419–425.

Andreu, C., del Olmo, M., 2020. Whole-Cell Biocatalysis in Seawater: New Halotolerant Yeast Strains for the Regio- and Stereoselectivity Reduction of 1-Phenylpropane-1,2-Dione in Saline-Rich Media. ChemBioChem. 21(11), 1621–1628. 10.1002/cbic.202000023.

Bai Flagfeldt, D., Siewers, V., Huang, L., Nielsen, J., 2009. Characterization of chromosomal integration sites for heterologous gene expression in *Saccharomyces cerevisiae*. Yeast. 26(10), 545–551. 10.1002/yea.1705.

Bamba, T., Inokuma, K., Hasunuma, T., Kondo, A., 2018. Enhanced cell-surface display of a heterologous protein using *SED1* anchoring system in *SED1*-disrupted *Saccharomyces cerevisiae* strain. J. Biosci. and Bioeng. 125(3), 306–310. 10.1016/j.jbiosc.2017.09.013.

Blomberg, A., 2022. Yeast osmoregulation – glycerol still in pole position. FEMS Yeast Res. 22(1), foac035. 10.1093/femsyr/foac035.

Breuer, U., Harms, H., 2006. *Debaryomyces hansenii* — an extremophilic yeast with biotechnological potential. Yeast. 23(6), 415–437. 10.1002/yea.1374.

Clipson, N.J.W., Jennings, D.H., Smith, J.L., 1989. The response to salinity at the microscopic level of the marine fungus *Dendryphiella salina* Nicot and Pugh as investigated stereologically. New Phytol. 113(1), 21–27. 10.1111/j.1469-8137.1989.tb02391.x.

Corrigan, M.W., Kerwin-Iosue, C.L., Kuczmarski, A.S., Amin, K.B., Wykoff, D.D., 2013. The Fate of Linear DNA in *Saccharomyces cerevisiae* and *Candida glabrata*: The Role of Homologous and Non-Homologous End Joining. PLOS ONE. 8(7), e69628. 10.1371/journal.pone.0069628.

Dague, E., Bitar, R., Ranchon, H., Durand, F., Yken, H.M., François, J.M., 2010. An atomic force microscopy analysis of yeast mutants defective in cell wall architecture. Yeast. 27(8), 673–684. 10.1002/yea.1801.

Dakal, T.C., Solieri, L., Giudici, P., 2014. Adaptive response and tolerance to sugar and salt stress in the food yeast *Zygosaccharomyces rouxii*. Int. J. Food Microbiol. 185, 140–157. 10.1016/j.ijfoodmicro.2014.05.015.

de la Fuente-Colmenares, I., González, J., Sánchez, N.S., Ochoa-Gutiérrez, D., Escobar-Sánchez, V., Segal-Kischinevzky, C., 2024. Regulation of Catalase Expression and Activity by DhHog1 in the Halotolerant Yeast *Debaryomyces hansenii* Under Saline and Oxidative Conditions. J. Fungi. 10(11), 740. 10.3390/jof10110740.

Donzella, S., Capusoni, C., Pellegrino, L., Compagno, C., 2021. Bioprocesses with Reduced Ecological Footprint by Marine *Debaryomyces hansenii* Strain for Potential Applications in Circular Economy. J. Fungi. 7(12), 1028. 10.3390/jof7121028.

Estrada, M., Navarrete, C., Møller, S., Procentese, A., Martínez, J.L., 2023a. Utilization of salt-rich by-products from the dairy industry as feedstock for recombinant protein production by *Debaryomyces hansenii*. Microb. Biotechnol. 16(2), 404–417. 10.1111/1751-7915.14179.

Estrada, M., Navarrete, C., Møller, S., Quirós, M., Martínez, J.L., 2023b. Open (non-sterile) cultivations of *Debaryomyces hansenii* for recombinant protein production combining industrial side-streams with high salt content. New Biotechnol. 78, 105–115. 10.1016/j.nbt.2023.10.005.

Fukuhara, H., 2006. Random AT library: autonomously replicating sequence (ARS) activity of chemically synthesized random sequences for transformation of nonconventional yeast species. FEMS Yeast Res. 6(8), 1281–1287. 10.1111/j.1567-1364.2006.00125.x.

Gui, Q., Deng, S., Zhou, Z., Cao, W., Zhang, X., Shi, W., Cai, X., Jiang, W., Cui, Z., Hu, Z., Chen, X., 2021. Transcriptome Analysis in Yeast Reveals the Externality of Position Effects. Mol. Biol. & Evol. 38(8), 3294–3307. 10.1093/molbev/msab104.

Gulati, S., Sun, Q., 2025. Complete Enzymatic Depolymerization of Polyethylene Terephthalate (PET) Plastic Using a *Saccharomyces cerevisiae*-Based Whole-Cell Biocatalyst. Environ. Sci. & Technol. Lett. 12(4), 419–424. 10.1021/acs.estlett.5c00190.

Hamada, T., Noda, F., Hayashi, K., 1984. Structure of Cell Wall and Extracellular Mannans from *Saccharomyces rouxii* and Their Relationship to a High Concentration of NaCl in the Growth Medium. Appl. & Environ. Microbiol. 48(4), 708–712. 10.1128/aem.48.4.708-712.1984.

Hosono, K., 1992. Effect of salt stress on lipid composition and membrane fluidity of the salt-tolerant yeast *Zygosaccharomyces rouxii*. Microbiology. 138(1), 91–96. 10.1099/00221287-138-1-91.

Innocente, N., Renoldi, N., Moret, E., Maifreni, M., Marino, M., 2023. Volatilome of brine-related microorganisms in a curd-based medium. J. Dairy Sci. 106(12), 8404–8414. 10.3168/jds.2022-23051.

Inokuma, K., Hasunuma, T., Kondo, A., 2014. Efficient yeast cell-surface display of exo- and endo-cellulase using the *SED1* anchoring region and its original promoter. Biotechnol. for Biofuels. 7(1), 8. 10.1186/1754-6834-7-8.

Inokuma, K., Kitada, Y., Bamba, T., Kobayashi, Y., Yukawa, T., den Haan, R., van Zyl, W.H., Kondo, A., Hasunuma, T., 2021. Improving the functionality of surface-engineered yeast cells by altering the cell wall morphology of the host strain. Appl. Microbiol. & Biotechnol. 105(14), 5895–5904. 10.1007/s00253-021-11440-6.

Kipnis, P., Thomas, N., Ovalle, R., Lipke, P.N., 2004. The ER-Golgi v-SNARE Bet1p is required for cross-linking α-agglutinin to the cell wall in yeast. Microbiology. 150(10), 3219–3228. 10.1099/mic.0.27189-0.

Kogej, T., Gorbushina, A.A., Gunde-Cimerman, N., 2006. Hypersaline conditions induce changes in cell-wall melanization and colony structure in a halophilic and a xerophilic black yeast species of the genus *Trimmatostroma*. Mycol. Res. 110(6), 713–724. 10.1016/j.mycres.2006.01.014.

Kralj Kunčič, M., Kogej, T., Drobne, D., Gunde-Cimerman, N., 2010. Morphological Response of the Halophilic Fungal Genus *Wallemia* to High Salinity. Appl. & Environ. Microbiol. 76(1), 329–337. 10.1128/AEM.02318-09.

Li, Y., Wang, X., Zhou, N.-Y., Ding, J., 2024. Yeast surface display technology: Mechanisms, applications, and perspectives. Biotechnol. Adv. 76, 108422. 10.1016/j.biotechadv.2024.108422.

Li, Z., Sharma, N.K., Weintraub, S., Young, E., 2025. Combinatorial metabolic engineering of alkane biosynthesis in the osmotolerant yeast *Debaryomyces hansenii* CBS 767. bioRxiv., 2025.2010.2023.684212. 10.1101/2025.10.23.684212.

Love, M.I., Huber, W., Anders, S., 2014. Moderated estimation of fold change and dispersion for RNA-seq data with DESeq2. Genome Biol. 15(12), 550. 10.1186/s13059-014-0550-8.

Lozančić, M., Žunar, B., Hrestak, D., Lopandić, K., Teparić, R., Mrša, V., 2021. Systematic Comparison of Cell Wall-Related Proteins of Different Yeasts. J. Fungi. 7(2), 128. 10.3390/jof7020128.

Maggi, R.G., Govind, N.S., 2004. Regulated expression of green fluorescent protein in *Debaryomyces hansenii*. J. Ind. Microbiol. and Biotechnol. 31(7), 301–310. 10.1007/s10295-004-0150-9.

Margesin, R., Schinner, F., 2001. Potential of halotolerant and halophilic microorganisms for biotechnology. Extremophiles. 5(2), 73–83. 10.1007/s007920100184.

Martinić Cezar, T., Lozančić, M., Novačić, A., Matičević, A., Matijević, D., Vallée, B., Mrša, V., Teparić, R., Žunar, B., 2023. Streamlining N-terminally anchored yeast surface display via structural insights into *S. cerevisiae* Pir proteins. Microb. Cell Fact. 22(1), 174. 10.1186/s12934-023-02183-2.

Martinić Cezar, T., Marđetko, N., Trontel, A., Paić, A., Slavica, A., Teparić, R., Žunar, B., 2024. Engineering *Saccharomyces cerevisiae* for the production of natural osmolyte glucosyl glycerol from sucrose and glycerol through Ccw12-based surface display of sucrose phosphorylase. J. Biol. Eng. 18(1), 69. 10.1186/s13036-024-00468-7.

Martinić Cezar, T., Paić, A., Prekpalaj, S., Teparić, R., Žunar, B., Stuparević, I., 2025. Measuring the capacity of yeast for surface display of cell wall-anchored protein isoforms by using β-lactamase as a reporter enzyme. FEBS Open Bio. 15(4), 573–579. 10.1002/2211-5463.13886.

Martinić Cezar, T., Paić, A., Žunar, B., Stuparević, I., Mrša, V., Teparić, R., 2026. Enhancing Yeast Surface Display: UPR, ERAD, and ER Dynamics in Recombinant Protein Production. Food Technol. & Biotechnol. 64(1), 4–19. 10.17113/ftb.64.01.26.8764.

Michán, C., Martínez, J.L., Alvarez, M.C., Turk, M., Sychrova, H., Ramos, J., 2013. Salt and oxidative stress tolerance in *Debaryomyces hansenii* and *Debaryomyces fabryi*. FEMS Yeast Res. 13(2), 180–188. 10.1111/1567-1364.12020.

Minhas, A., Biswas, D., Mondal, A.K., 2009. Development of host and vector for high-efficiency transformation and gene disruption in *Debaryomyces hansenii*. FEMS Yeast Res. 9(1), 95–102. 10.1111/j.1567-1364.2008.00457.x.

Morais, A.R.C., Duarte, L.C., Lourenço, P., Torrado, I., Brás, T., Neves, L.A., Carvalheiro, F., 2023. Xylitol Production by *Debaryomyces hansenii* in Extracted Olive Pomace Dilute-Acid Hydrolysate. Fermentation. 9(12), 1020. 10.3390/fermentation9121020.

Navarrete, C., Estrada, M., Martínez, J.L., 2022a. Debaryomyces hansenii: an old acquaintance for a fresh start in the era of the green biotechnology. World J. Microbiol. & Biotechnol. 38(6), 99. 10.1007/s11274-022-03280-x.

Navarrete, C., Frost, A.T., Ramos-Moreno, L., Krum, M.R., Martínez, J.L., 2021. A physiological characterization in controlled bioreactors reveals a novel survival strategy for *Debaryomyces hansenii* at high salinity. Yeast. 38(5), 302–315. 10.1002/yea.3544.

Navarrete, C., Sánchez, B.J., Savickas, S., Martínez, J.L., 2022b. DebaryOmics: an integrative –omics study to understand the halophilic behaviour of *Debaryomyces hansenii*. Microb. Biotechnol. 15(4), 1133–1151. 10.1111/1751-7915.13954.

Navarrete, C., Siles, A., Martínez, J.L., Calero, F., Ramos, J., 2009. Oxidative stress sensitivity in *Debaryomyces hansenii*. FEMS Yeast Res. 9(4), 582–590. 10.1111/j.1567-1364.2009.00500.x.

Norkrans, B., 1966. Studies on marine occurring yeasts: Growth related to pH, NaCl concentration and temperature. Arch. für Mikrobiol. 54(4), 374–392. 10.1007/BF00406719.

Novačić, A., Beauvais, V., Oskomić, M., Štrbac, L., Dantec Aurélia, L., Rahmouni, A.R., Stuparević, I., Weis, K., 2021. Yeast RNA exosome activity is necessary for maintaining cell wall stability through proper protein glycosylation. Mol. Biol. Cell. 32(5), 363–375. 10.1091/mbc.E20-08-0544-T.

O’Riordan, N.M., Jurić, V., O’Neill, S.K., Roche, A.P., Young, P.W., 2024. A Yeast Modular Cloning (MoClo) Toolkit Expansion for Optimization of Heterologous Protein Secretion and Surface Display in *Saccharomyces cerevisiae*. ACS Synth. Biol. 13(4), 1246–1258. 10.1021/acssynbio.3c00743.

Ochoa-Gutiérrez, D., Reyes-Torres, A.M., de la Fuente-Colmenares, I., Escobar-Sánchez, V., González, J., Ortiz-Hernández, R., Torres-Ramírez, N., Segal-Kischinevzky, C., 2022. Alternative CUG Codon Usage in the Halotolerant Yeast *Debaryomyces hansenii*: Gene Expression Profiles Provide New Insights into Ambiguous Translation. J. Fungi. 8(9), 970. 10.3390/jof8090970.

Přibylová, L., Farkaš, V., Slaninová, I., de Montigny, J., Sychrová, H., 2007. Differences in osmotolerant and cell-wall properties of two *Zygosaccharomyces rouxii* strains. Folia Microbiol. 52(3), 241–245. 10.1007/BF02931305.

Prista, C., Almagro, A., Loureiro-Dias, M.C., Ramos, J., 1997. Physiological basis for the high salt tolerance of *Debaryomyces hansenii*. Appl. & Environ. Microbiol. 63(10), 4005–4009. 10.1128/aem.63.10.4005-4009.1997.

Prista, C., Michán, C., Miranda, I.M., Ramos, J., 2016. The halotolerant *Debaryomyces hansenii*, the Cinderella of non-conventional yeasts. Yeast. 33(10), 523–533. 10.1002/yea.3177.

Ragni, E., Piberger, H., Neupert, C., García-Cantalejo, J., Popolo, L., Arroyo, J., Aebi, M., Strahl, S., 2011. The genetic interaction network of *CCW12*, a *Saccharomyces cerevisiae* gene required for cell wall integrity during budding and formation of mating projections. BMC Genom. 12(1), 107. 10.1186/1471-2164-12-107.

Ragni, E., Sipiczki, M., Strahl, S., 2007. Characterization of Ccw12p, a major key player in cell wall stability of *Saccharomyces cerevisiae*. Yeast. 24(4), 309–319. 10.1002/yea.1465.

Rajkumar, A.S., Liu, G., Bergenholm, D., Arsovska, D., Kristensen, M., Nielsen, J., Jensen, M.K., Keasling, J.D., 2016. Engineering of synthetic, stress-responsive yeast promoters. Nucleic Acids Res. 44(17), e136–e136. 10.1093/nar/gkw553.

Ramos-Moreno, L., Ramos, J., Michán, C., 2019. Overlapping responses between salt and oxidative stress in *Debaryomyces hansenii*. World J. Microbiol. & Biotechnol. 35(11), 170. 10.1007/s11274-019-2753-3.

Ribeiro, R.A., Vitorino, M.V., Godinho, C.P., Bourbon-Melo, N., Robalo, T.T., Fernandes, F., Rodrigues, M.S., Sá-Correia, I., 2021. Yeast adaptive response to acetic acid stress involves structural alterations and increased stiffness of the cell wall. Sci. Rep. 11(1), 12652. 10.1038/s41598-021-92069-3.

Robinson, D., Vanacloig-Pedros, E., Cai, R., Place, M., Hose, J., Gasch, A.P., 2023. Gene-by-environment interactions influence the fitness cost of gene copy-number variation in yeast. G3 Genes|Genomes|Genet. 13(10), jkad159. 10.1093/g3journal/jkad159.

Rodríguez-Peña, J.M., García, R., Nombela, C., Arroyo, J., 2010. The high-osmolarity glycerol (HOG) and cell wall integrity (CWI) signalling pathways interplay: a yeast dialogue between MAPK routes. Yeast. 27(8), 495–502. 10.1002/yea.1792.

Ruiz-Pérez, F.S., Ruiz-Castilla, F.J., Leal, C., Martínez, J.L., Ramos, J., 2023. Sodium and lithium exert differential effects on the central carbon metabolism of *Debaryomyces hansenii* through the glyoxylate shunt regulation. Yeast. 40(7), 265–275. 10.1002/yea.3856.

Sánchez, B.J., 2021. debary-omics: Analysis of omics data for *Debaryomyces hansenii*. GitHub. https://github.com/computer-aided-biotech/debary-omics.

Sánchez, N.S., Calahorra, M., González-Hernández, J.C., Peña, A., 2006. Glycolytic sequence and respiration of *Debaryomyces hansenii* as compared to *Saccharomyces cerevisiae*. Yeast. 23(5), 361–374. 10.1002/yea.1360.

Sánchez, N.S., Calahorra, M., González, J., Defosse, T., Papon, N., Peña, A., Coria, R., 2020. Contribution of the mitogen-activated protein kinase Hog1 to the halotolerance of the marine yeast *Debaryomyces hansenii*. Curr. Genet. 66(6), 1135–1153. 10.1007/s00294-020-01099-3.

Serra-Cardona, A., Petrezsélyová, S., Canadell, D., Ramos, J., Ariño, J., 2014. Coregulated Expression of the Na+/Phosphate Pho89 Transporter and Ena1 Na+-ATPase Allows Their Functional Coupling under High-pH Stress. Mol. & Cell. Biol. 34(24), 4420–4435. 10.1128/MCB.01089-14.

Serrano, R., Ruiz, A., Bernal, D., Chambers, J.R., Ariño, J., 2002. The transcriptional response to alkaline pH in *Saccharomyces cerevisiae*: evidence for calcium-mediated signalling. Mol. Microbiol. 46(5), 1319–1333. 10.1046/j.1365-2958.2002.03246.x.

Sharma, P., Meena, N., Aggarwal, M., Mondal, A.K., 2005. *Debaryomyces hansenii*, a highly osmo-tolerant and halo-tolerant yeast, maintains activated Dhog1p in the cytoplasm during its growth under severe osmotic stress. Curr. Genet. 48(3), 162–170. 10.1007/s00294-005-0010-9.

Šupljika, N., Paić, A., Novačić, A., Martinić Cezar, T., Vallée, B., Teparić, R., Stuparević, I., Žunar, B., 2025. *Saccharomyces cerevisiae* Mub1, a substrate adaptor of E3 ubiquitin ligase Ubr2, modulates sensitivity to cell wall stressors through multiple transcription factors. The FEBS J. 292(19), 5046–5066. 10.1111/febs.70091.

Thomé, P.E., 2007. Cell wall involvement in the glycerol response to high osmolarity in the halotolerant yeast *Debaryomyces hansenii*. Antonie van Leeuwenhoek. 91(3), 229–235. 10.1007/s10482-006-9112-8.

Turk, M., Montiel, V., Žigon, D., Plemenitaš, A., Ramos, J., 2007. Plasma membrane composition of *Debaryomyces hansenii* adapts to changes in pH and external salinity. Microbiology. 153(10), 3586–3592. 10.1099/mic.0.2007/009563-0.

Villarreal-Huerta, D., Mendoza-Téllez, B., Rosas-Paz, M.Á.A., Sánchez, N.S.S., Jiménez-Nava, R.A.A., Cristiani-Urbina, E., Segal-Kischinevzky, C., González, J., 2026. Hog1 MAP kinase modulates early riboflavin accumulation under low-pH and saline conditions in *Debaryomyces hansenii*. Front. in Microbiol. Volume 17–2026. 10.3389/fmicb.2026.1746023.

Wang, D., Zhang, M., Huang, J., Zhou, R., Jin, Y., Wu, C., 2020. *Zygosaccharomyces rouxii* Combats Salt Stress by Maintaining Cell Membrane Structure and Functionality. J. Microbiol. Biotechnol. 30(1), 62–70. 10.4014/jmb.1904.04006.

Wang, Y., Deng, Q., Liu, F., Li, Q., Li, R., Wang, H., Sun, C., Wang, X., Wu, X., Chen, Y., 2024. Urea amidolyase displayed on the surface of *Saccharomyces cerevisiae* for ethyl carbamate reduction in Chinese Huangjiu. Food Control. 166, 110685. 10.1016/j.foodcont.2024.110685.

Wu, X.-L., Li, B.-Z., Zhang, W.-Z., Song, K., Qi, H., Dai, J.-b., Yuan, Y.-J., 2017. Genome-wide landscape of position effects on heterogeneous gene expression in *Saccharomyces cerevisiae*. Biotechnol. for Biofuels. 10(1), 189. 10.1186/s13068-017-0872-3.

Yang, X., Xiao, S., Wang, J., 2024. *Debaryomyces hansenii* Strains from Traditional Chinese Dry-Cured Ham as Good Aroma Enhancers in Fermented Sausage. Fermentation. 10(3), 152. 10.3390/fermentation10030152.

Yap, W.Y., Schiestl, R.H., 1995. Nature of abortive transformation in *Saccharomyces cerevisiae*. Curr. Genet. 28(6), 517–520. 10.1007/BF00518163.

Zhu, A., Ibrahim, J.G., Love, M.I., 2019. Heavy-tailed prior distributions for sequence count data: removing the noise and preserving large differences. Bioinformatics. 35(12), 2084–2092. 10.1093/bioinformatics/bty895.

